# Exposure time to moderately high temperature affects photosynthetic response to water deficit in *Eucalyptus globulus* Labill

**DOI:** 10.1101/2025.05.16.654555

**Authors:** Maria Teresa Portes, Daniel S.C. Damineli, Maria Manuela Chaves, Gustavo M. Souza

## Abstract

The co–occurrence of drought and heat stress in the field is especially frequent and the plant response to a combination of these abiotic stresses is singular, remaining largely unknown. Understanding physiological responses to a combined stress is of particular importance in face of the imminent global climate changes, which is predicted to entail not only higher temperatures, but also increase in drought, imposing restrictions to the productivity of economically important species as *E. globulus*. This forest species is critically affected by changes in temperature and water availability, since its high productivity implies in high rates of water use. In order to evaluate the photosynthetic responses of *Eucalyptus globulus* to these environmental conditions, young plants were grown at 25° C and 35° C under different irrigation regimes: full irrigation – FI, moderate drought – MD, and severe drought – SD. Plants grown at 25° C FI and MD were also subjected to a short–term exposure to 35° C. Several ecophysiological variables, derived mainly from leaf gas exchange and chlorophyll *a* fluorescence measurements, were compared in order to test the hypotheses that: *i) E. globulus* show different responses to short or long–term exposure to 35° C; *ii)* the effect of both stresses combined will not be simply the sum of each condition applied separately. Our results showed that plants respond differently to water deficit according to the thermal regime. The combination of water deficit and a long–term exposure to 35° C caused a down–acclimation of the photosynthetic capacity, while plants under the short–term treatment showed a higher performance, even when compared to the 25° C control condition, supporting the hypothesis *i*. Thus, the short–term exposure to 35° C induced an increase in tolerance to water deficit. Since the response to a combination of heat and drought differed from the superposition of individual responses, as observed mainly at 35° C, the hypothesis *ii* was also supported by our results.

## INTRODUCTION

Generally, in the field, several abiotic stresses occur simultaneously, being especially frequent the co–occurrence of drought and heat stress. Increase in leaf temperature is also a consequence of drought since plants decrease transpirational cooling due to stomatal closure (SHARKEY, 2005). Although both stress types are associated, molecular and metabolic studies suggest that the plant response to a combination of drought and heat is unique and cannot be directly extrapolated from the individual response to each stress (RIZHSKY et al., 2002, 2004).

Given that photosynthesis is a central process to cellular metabolism, involving large fluxes of carbon, nitrogen and energy (LAWLOR, 2001) and being integrated with respiration, electron transport and ATP synthesis in the mitochondria (ATKIN & MACHEREL, 2009), it is considerably vulnerable to water deficit and temperature. Indeed photosynthesis, together with cell growth, is among the main processes to be affected by drought (CHAVES, 1991). Direct effects are principally the decreased CO_2_ availability caused by diffusion limitations through the stomata and mesophyll (CHAVES et al., 2003, 2009; FLEXAS et al., 2004, 2007) and alterations in the photosynthetic metabolism (LAWLOR & CORNIC, 2002; LAWLOR & TEZARA, 2009). Moreover, under drought, the balance between energy harvesting and metabolism may be disturbed, leading to a decrease in the efficiency of the photochemical phase of photosynthesis, occurring in parallel with an increase in energy dissipation in the chloroplast (DEMMIG–ADAMS & ADAMS, 1992). Additionally, when drought becomes severe oxidative stress may arise due to the generation of reactive oxygen species (REDDY et al., 2004).

Regarding the effects of high temperature, photochemical reactions in the thylakoid lamellae and carbon metabolism in the stroma of the chloroplasts have been suggested to be the primary sites of injury (WISE et al., 2004). Direct injuries due to high temperatures include protein denaturation and aggregation, as well as an increase in the fluidity of membrane lipids and electrolyte leakage (WAHID et al., 2007). Plant metabolism can undergo acclimation after a prolonged exposure to a different growth temperature, which could eventually result in metabolic homeostasis, for example, with the maintenance of similar photosynthetic and respiratory rates (STITT & HURRY, 2002; ATKIN & TJOELKER, 2003). Furthermore, an indication of thermal acclimation is the deviation of the temperature optimum towards the new growth temperature (BERRY & BJÖRKMAN, 1980), as reported in *Eucalyptus globulus* by BATTAGLIA et al. (1996).

Thus far, most studies on plant physiology have focused on the impact of a single environmental stress, *e.g.* either water deficit or heat shock (REDDY et al., 2004; CAMEJO et al., 2005), whereas the combination of stress types has received less attention (RIZHSKY et al., 2004). The effects of the combination of drought and heat stress were previously studied on the growth and productivity of maize, barley, sorghum and different grasses (MITTLER, 2006) but, to our knowledge, there was no attempt to evaluate the combined effect of water stress and moderately high temperature in tree species, as *Eucalyptus globulus*. *E. globulus* is a forest species with considerable economical importance, once it is the foremost pulpwood eucalypt species planted in temperate regions (DOUGHTY, 2000) and its productivity is particularly sensitive to drought, given its high water demand (WHITEHEAD & BEADLE, 2004).

Understanding how this plant species respond to increases in water deficit and elevated temperature is particularly important in face of the imminent global climate changes, given that the predicted global warming, in the next century, by an average of 2–4° C in conjunction with changes in precipitation will likely lead to droughts and supra–optimal temperatures in many areas of the globe (IPCC, 2007). Drought may induce large–scale declines in tree growth in temperate forests, once the productivity of forest ecosystems is severely constrained by water availability (BRÉDA et al., 2006). In addition, temperature–mediated changes in leaf photosynthesis and respiration are now accepted as important components of the biosphere’s response to global climate change (ATKIN & TJOELKER, 2003). Thus, the present study has the aim to evaluate the effects of water deficit on the photosynthetic capacity in plants of *E. globulus* under different thermal regimes.

In order to evaluate the photosynthetic responses of *E. globulus* to these environmental conditions, young plants were grown at 25° C and 35° C under different irrigation regimes: full irrigation – FI, moderate drought – MD, and severe drought – SD. Plants grown at 25° C FI and MD were also subjected to a short–term exposure to 35° C. Several ecophysiological variables, derived mainly from leaf gas exchange and chlorophyll *a* fluorescence measurements, were compared in order to test the hypotheses that: *i) E. globulus* show different responses to short or long–term exposure to 35° C; *ii)* the effect of both stresses combined will not be simply the sum of each condition applied separately (MITTLER, 2006). We will show a marked difference between the responses to long and short–term exposure to 35° C, in different irrigation treatments, where the short–term treatment induced a lower down–acclimation of the photosynthetic capacity than the long–term and, furthermore, promoted a potential increase in tolerance to water deficit.

## MATERIAL AND METHODS

### Plant material and growth conditions

Seeds of *Eucalyptus globulus* Labill. were germinated in pots in a fully sunlit greenhouse where they remained for about 1 month. Seedlings with one or two pairs of true leaves were then transferred to 3 L plastic pots containing a 2:1 (v/v) soil and sand mixture and placed in a controlled growth chamber (FITOCLIMA 10000EHHF, Aralab) set to maintain temperature at 25/18° C (day/night cycle), 800 μmol m^-2^ s^-1^ of photosynthetic photon flux density (PPFD) provided with fluorescent tubes, 60% of relative humidity and a photoperiod of 16 h. Healthy 5 months–old plants were then transferred to another controlled growth chamber (FITOCLIMA 700EDTU, Aralab) with identical conditions, where they remained for 1 month to ensure acclimation before experiments started. Until the beginning of the experiments, pots were irrigated daily to full soil capacity and a standard Hoagland’s solution was applied every ten days in order to prevent deficiency in essential nutrients.

### Experimental set up

Experiments were performed in leaves grown under two diurnal thermal conditions: *i)* 25/18° C, and *ii)* 35/18° C (day/night cycle). In each thermal treatment, six plants were randomly selected for three irrigation treatments: *i)* irrigated to field capacity (full irrigation – FI), *ii)* irrigated to 25% of the field capacity (moderate drought treatment – MD), and *iii)* irrigated to 12.5% of the field capacity (severe drought treatment – SD). Measurements were performed two weeks after the beginning of the irrigation treatments, and pots were frequently rotated to minimize microclimatic effects inside the growth chamber. Additionally, a heat shock experiment was performed, consisting in exposing plants grown at 25° C to an abrupt increase in diurnal temperature to 35° C, remaining under such condition for four days. This treatment, designated “35° C Short– term”, was performed under FI and MD conditions where the water deficit was applied one week before the heat shock.

### Leaf gas exchange and chlorophyll a fluorescence measurements

Light (*A*_N_–PPFD) and CO_2_ (*A*_N_–C_i_) response curves were performed, where leaf gas exchange parameters were determined simultaneously with chlorophyll fluorescence measurements using an open infrared gas–exchange analyzer (Li–6400, Li–Cor Inc., Lincoln, NE, USA) with an integrated fluorescence chamber head (Li–6400-40; Li–Cor Inc.) that provides LED–based fluorescence and irradiation. Measurements were taken between 8:00–12:00 h in young fully expanded leaves from the light exposed parts of the shoots plants acclimated to the above mentioned conditions. Measurements were performed with a 2 cm^2^ Li–6400 leaf chamber in one attached leaf in each of the six plants evaluated per irrigation and thermal treatment. Net CO_2_ assimilation (*A*_N_ – μmol CO_2_ m^-2^ s^-1^) and transpiration (E – mmol H_2_O m^-2^ s^-1^) rates, stomatal conductance (*g*_s_ – mol H_2_O m^-2^ s^-1^) and intercellular CO_2_ concentration (C_i_ – μmol CO_2_ mol^-1^) were calculated using the LI–6400 data analysis program which uses VON CAEMMERER & FARQUHAR’s (1981) general gas exchange formula. Environmental conditions in the leaf chambers were controlled with the LI–6400, maintaining conditions practically identical to the growth chamber conditions (described above). During all measurements vapour pressure deficit (VPD) was maintained around 1.0–1.5 kPa using a dew point generator (Li–610, Licor) attached to the LI–6400.

Light response curves (*A*_N_–PPFD) were obtained by gradually varying PPFD from 1400 to 25 μmol photon m^-2^ s^-1^ (in ten levels) under a constant CO_2_ concentration of 380 μmol mol^−1^ and block temperature at 25° C or 35° C, according to thermal treatment. PPFD was provided by the Li– 6400–40 light source, with 10% blue light to maximize stomatal aperture. Measurements were recorded with 4–5 min intervals between each reading and were logged when the total coefficient of variation (CV) was ≤1 %, indicating stability of the reading. CO_2_ response curves (*A*_N_–C_i_) were performed at 1000 μmol photon m^-2^ s^-1^ of PPFD with CO_2_ concentration varying from 100 to 1500 μmol mol^-1^. Measurements were also recorded when *A* values were stable (CV ≤1%), 4–5 min after each change in chamber CO_2_ concentration.

### Model and derived parameters

Light and CO_2_ response curves were fitted using the following equation (FARQUHAR et al., 1980)

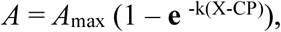

where *A* is the net CO_2_ assimilation, *A*_max_ is the maximum CO_2_ assimilation, **e** is the Euler’s number, k is a constant related to the convexity of the curve, X is PPFD or C_i_ and CP is the light (LCP) or CO_2_ compensation point (CCP). Light and CO_2_ saturation points, LSP and CSP respectively, were estimated calculating the values in which *A* reached 90% of *A*_max_, while apparent quantum (α) and carboxylation (ε) efficiencies were estimated using the initial linear slope of *A*_N_– PPFD and *A*_N_*–*C_i_ curves respectively.

Relative stomatal limitation of photosynthesis (L_s_) was calculated from *A*_N_*–*C_i_ curves as proposed by FARQUHAR & SHARKEY (1982):

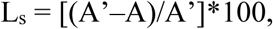

where A’ is the CO_2_ assimilation when C_i_ equals the atmospheric concentration (380 μmol mol^−1^) and A is the CO_2_ assimilation when CO_2_ concentration in the sample chamber (C_e_) equals the atmospheric concentration. The maximum rate of Rubisco carboxylation (*V*_cmax_ – μmol CO_2_ m^-2^ s^-1^) and maximum electron transport rate at saturating light (*J*_max_ – μmol CO_2_ m^-2^ s^-1^) were determined on a C_i_ basis from *A*_N_–C_i_ curves by fitting the model of FARQUHAR et al. (1980) with modifications by SHARKEY (1985) to *A*_N_ –C_i_ response curves, as described by MAROCO et al. (2002).

Mesophyll conductance (*g*_m_ – mol CO_2_ m^-2^ s^-1^ bar^-1^) was estimated with the method of HARLEY et al. (1992) using the following equation:

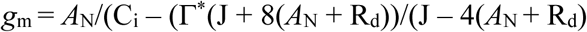

where J is the photosynthetic electron transport rate and Γ* is the CO_2_ compensation point in the absence of R_d_, which is the mitochondrial respiration occurring during the day, estimated using the LAISK (1977) method. The mean values of *g*_m_ presented correspond to C_e_ concentration around 380 μmol mol^-1^.

Intrinsic water use efficiency (WUE_i_ – μmol CO_2_ mol H_2_O^-1^) was calculated with *A*_N_/*g*_s_ ratio from *A*_N_–PPFD response curves. Leaf dark respiration (R_d_ – μmol CO_2_ m^-2^ s^-1^) was calculated with *A* values at PPFD = 0 measured in light response curves of leaves adapted to 30 min of darkness.

### Chlorophyll a fluorescence parameters

Maximal (F_M_) and basal (F_0_) fluorescence yields were measured in the same dark–adapted leaves used to estimated R_d_, whereas steady–state (F_S_) and maximal (F’_M_) fluorescence were measured in light–adapted leaves during the *A*_N_–PPFD curves (VAN KOOTEN & SNEL, 1990). Thus, variable fluorescence yield was determined in both dark– adapted (F_V_= F_M_ – F_0_) and light–adapted (ΔF= F’_M_ – F_S_) states. The fluorescence parameters calculated were: potential (F_V_/F_M_) and effective (<!PSII= ΔF/F’_M_) quantum efficiency of the photosystem II (PSII), photochemical {qP= [(F’_M_ – F_S_)/(F’_M_ – F’_0_)]} and non photochemical {NPQ= [(F_M_ – F’_M_)/F’_M_]} fluorescence quenching, PSII antennae efficiency {F’_V_/F’_M_= [(F’_M_–F_0_)/ F’_M_]}, and apparent electron transport rate [ETR= (PPFD x (ΔF/F’_M_ x 0.5 x 0.84)] (BILGER et al., 1995; MAXWELL & JOHNSON, 2000). F’_M_ was obtained with a pulse of saturating actinic light applied simultaneously to the gas exchange measurements, where F_S_ is the steady-state fluorescence measured briefly before the saturating light pulse allowing estimation of the PSII quantum yield (GENTY et al., 1989). F_0_’ is the basal fluorescence yield after photosystem I excitation by far–red light. For the calculation of ETR, 0.5 was used as the fraction of excitation energy distributed to PSII, and 0.84 as the fraction of light absorption (DEMMIG & BJÖRKMAN, 1987).

### Leaf water potential (Ψ_w_)

Pre–dawn (Ψ_pd_) and midday (Ψ_md_) leaf water potential were measured at 5:00 and 13:00 h, respectively, with a Scholander–type pressure chamber (PMS Instruments Co., Corvallis, OR, USA) in six fully expanded leaves of six different plants per treatment (n=6). The leaf water potential was recorded in the same days where gas exchange measurements were performed.

### Statistical analysis

The experiment was arranged in a randomized design with 6 replicates. All data were subjected to a two–way analyses of variance (ANOVA) followed by a post–hoc Tukey’s test at the 0.05 significance level in order to assess the effects of thermal and irrigation treatments and their interaction on the different dependent variables evaluated. Statistical data analyses were performed with Statistica (v7, Statsoft Tulsa, OK, USA). Throughout the text every difference considered significant had a *p* < 0.05 (Tukey’s test).

Principal component analysis (PCA) is a technique which enables the exploration of multivariate data sets through the reduction of n variables to lower dimensions, which are formed by principal components. These are independent axes composed of linear combinations of the original variables, as to maximize the explained variances. Thus, the first principal component (PC1) explains most of the variance in the observed data, followed by PC2 and so on, being an effective approach to identify groups formed by the combined effect of the evaluated variables (HAIR et al., 2006). The analysis was performed in the language for statistical computing R, by means of the function *prcomp* (R Development Core Team, 2009), on the normalized data of all replicates (variances scaled to 1) with the variables: C_i_, *A*_N_, *g*_s_, *g*_m_, *V*_cmax_, *J*_max_, F_0_, F_M_, F_V_/F_M_, F_V_/F_0_, L_s_, R_d_, *A*_max light_, LCP, α, LSP, *A*_max CO_2, CCP, ε, CSP and WUE_i_.

## RESULTS

The reduction in water availability resulted in a decrease in leaf water potential (Ψ_w_) according to drought severity, both in predawn (Ψ_pd_) and midday (Ψ_md_), being significantly the lowest under SD at 25° C and 35° C and in the MD at 35° C Short–term (Figure 1). The decrease in Ψ_w_ and the increase in temperature constrained the photosynthetic apparatus of *E. globulus*, an effect derived from photochemical, biochemical and CO_2_ diffusion limitations.

**Figure 1.**
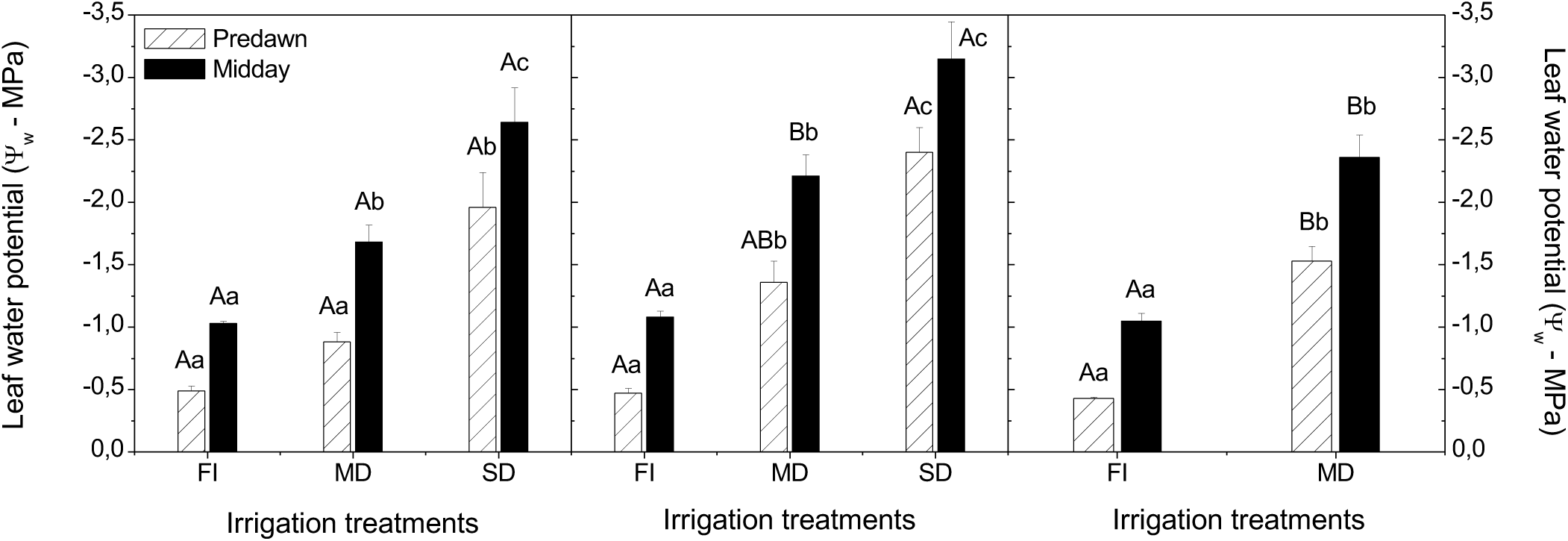
Leaf water potential (Ψ_w_) at predawn (Ψ_pd_) and midday (Ψ_md_) measured in *Eucalyptus globulus* grown at 25° C, 35° C and 35° C Short-term under different irrigation treatments: full irrigation (FI), moderate drought (MD) and severe drought (SD). Data represent means ± SE (n=6).

### Photochemical limitations

The effects of water deficit in the photochemical apparatus were different in the distinct thermal treatments, as evidenced by the fluorescence parameters measured in light and dark– adapted leaves (Table 1, Figure 2). Fluorescence parameters in dark–adapted leaves indicated that the long–term exposure to 35° C disturbed the photochemical apparatus mainly in the response to water deficit, given the significantly higher increase in basal (F_0_) and maximal (F_M_) fluorescence yields observed from FI to MD and SD. Curiously, at 35° C Short–term, F_0_ decreased under MD and F_M_ was similar under both irrigation regimes, indicating that the photochemical apparatus was practically unaffected by a short exposure to 35° C. The potential quantum efficiency (F_V_/F_M_) and the variable to basal fluorescence ratio (F_V_/F_0_) generally decreased with reduced water availability, showing a prominent trend to decrease at 35° C, whereas at 35° C Short–term both parameters showed a non–significant increase under MD (Table 1).

**Figure 2.**
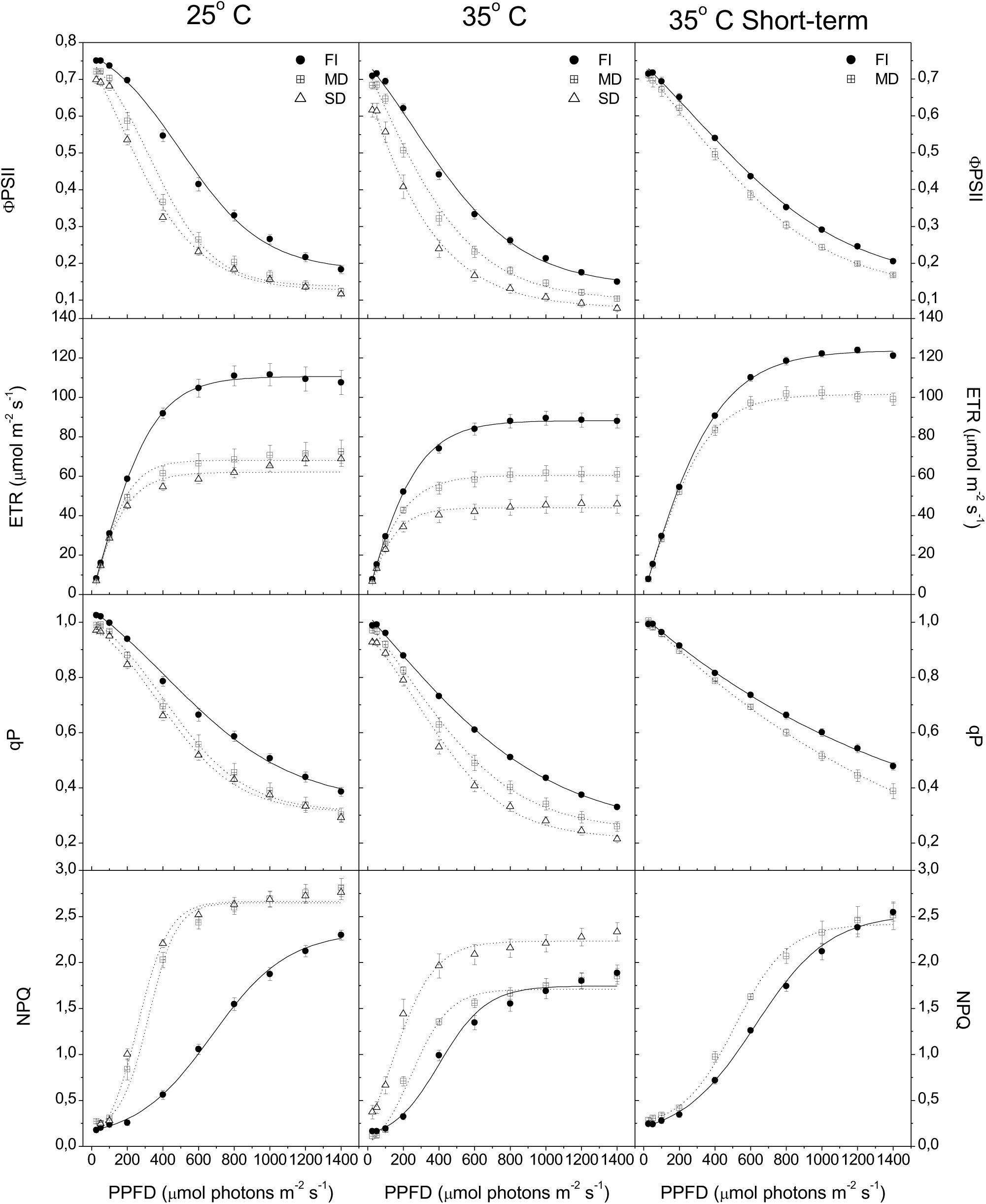
Effective quantum efficiency of PSII (<λPSII), electron transport rate (ETR), photochemical quenching (qP) and non-photochemical quenching (NPQ) light response curves in leaves of *Eucalyptus*

**Table 1.** Chlorophyll *a* fluorescence parameters in dark–adapted leaves of *Eucalyptus globulus* grown at 25° C, 35° C and 35° C Short-term and exposed to different irrigation regimes: full irrigation (FI), moderate drought (MD) and severe drought (SD). Data represent means ± SE (n=6). Different capital letters express significant difference for thermal treatments and different lower-case letters express significant differences for irrigation treatments (Tukey’s test, *p* < 0.05). F_0_ = basal fluorescence yield, F_M_ = maximal fluorescence yield, F_V_/F_M_ = potential quantum efficiency, F_V_/F_0_ = variable to basal fluorescence ratio.

| Parameters | 25° C |  |  | 35° C |  |  | 35° C Short-term |  |
| --- | --- | --- | --- | --- | --- | --- | --- | --- |
|  | FI | MD | SD | FI | MD | SD | FI | MD |
| $F_0$ | 398.47 ± 11.52 <sup>Aa</sup> | 416.95 ± 9.12 <sup>Ba</sup> | 435.88 ± 17.13 <sup>Ba</sup> | 437.70 ± 12.24 <sup>Ab</sup> | 565.75 ± 52.89 <sup>Aab</sup> | 651.32 ± 78.04 <sup>Aa</sup> | 475.3 ± 18.78 <sup>Aa</sup> | 451.1 ± 15.68 <sup>ABa</sup> |
| $F_M$ | 2120.35 ± 28.73 <sup>Aa</sup> | 2177.55 ± 33.51 <sup>Aa</sup> | 2149.34 ± 47.98 <sup>Aa</sup> | 2020.75 ± 35.27 <sup>Aa</sup> | 2211.97 ± 62.47 <sup>Aab</sup> | 2325.47 ± 100.42 <sup>Ab</sup> | 2120.58 ± 11.83 <sup>Aa</sup> | 2124.48 ± 29.8 <sup>Aa</sup> |
| $F_V/F_M$ | 0.812 ± 0.01 <sup>Aa</sup> | 0.808 ± 0.01 <sup>Aa</sup> | 0.797 ± 0.01 <sup>Aa</sup> | 0.783 ± 0.01 <sup>Aa</sup> | 0.745 ± 0.02 <sup>Aa</sup> | 0.715 ± 0.04 <sup>Bb</sup> | 0.776 ± 0.01 <sup>Aa</sup> | 0.788 ± 0.01 <sup>Aa</sup> |
| $F_V/F_0$ | 4.352 ± 0.22 <sup>Aa</sup> | 4.230 ± 0.10 <sup>Aa</sup> | 3.951 ± 0.15 <sup>Aa</sup> | 3.628 ± 0.11 <sup>Aa</sup> | 3.037 ± 0.31 <sup>Aa</sup> | 2.759 ± 0.54 <sup>Aa</sup> | 3.49 ± 0.18 <sup>Aa</sup> | 3.725 ± 0.16 <sup>Aa</sup> |

Fluorescence parameters in light–adapted leaves were measured concomitantly with light response curves (*A*_N_–PPFD), thus their courses are presented as a function of PPFD (Figure 2). Effective quantum efficiency (ΦPSII) and photochemical quenching (qP) showed similar courses, both being considerably reduced under MD and SD at 25° C and 35° C, whereas a lower decrease due to water deficit is verifiable at 35° C Short–term. Electron transport rate (ETR) was more affected by the long–term exposition to 35° C since it markedly decreased from FI to MD and SD, when compared to 25° C. Water deficit had a remarkably lower effect in ETR at 35° C Short–term, even when compared to 25° C, showing the highest values under MD. Curiously, non– photochemical quenching (NPQ) was higher at 25° C than at 35° C in all irrigation treatments, also showing a pronounced difference between water deficit and FI treatments which was not verifiable in both 35° C treatments (Figure 2).

### Biochemical and CO_2_ diffusion limitations

The exposure time to 35° C lead to different photosynthetic responses in fully irrigated plants given that, comparing to 25° C FI, *A*_max light_ was significantly lower at 35° C while at 35° C Short–term no significant difference was detected (Table 2). Such response can also be verified through the *A*_N_–PPFD curves, which showed similar courses in plants under FI at 25° C and 35° C Short–term, while at 35° C the CO_2_ assimilation course was lower possibly related with the lower stomatal conductance (*g*_s_) observed in this treatment (Figure 3). Interestingly, the effects of water deficit on the photosynthetic efficiency were significantly less pronounced at 35° C Short–term, showing higher *A*_max light_ under MD compared to the other thermal treatments (Table 2). Values of *A*_max light_ are clearly related with their respective *g*_s_, as can be seen in the reduction in *A*_N_ from FI to water deficit treatments, between 25° C and 35° C under FI, and in the maintenance of a higher *A*_N_ and *g*_s_ at 35° C Short–term under MD (Figure 3).

**Figure 3.**
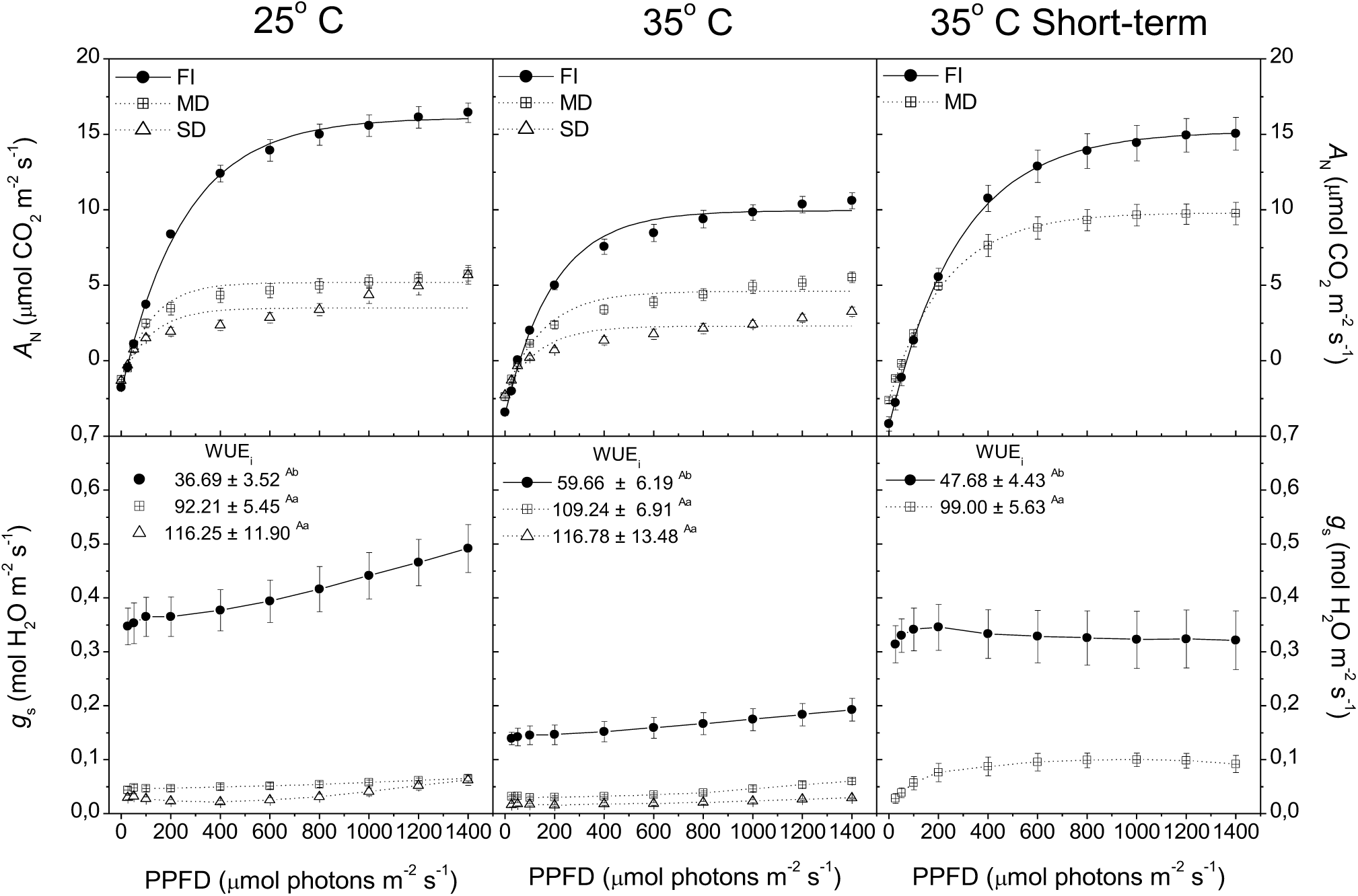
Net CO_2_ assimilation (*A*_N_) and stomatal conductance (*g*_s_) light response curves, together with intrinsic water use efficiency (WUE_i_ – μmol CO_2_ mol H_2_O^-1^) in leaves of *Eucalyptus globulus* grown at 25° C, 35° C and 35° C Short-term under different irrigation treatments: full irrigation (FI), moderate drought (MD) and severe drought (SD). PPFD is the photosynthetic photon flux density. Data represent means ± SE (n=6). Different capital letters express significant differences among thermal treatments and different lower-case letters express significant differences among irrigation treatments (Tukey’s test, *p* < 0.05).

**Table 2.**
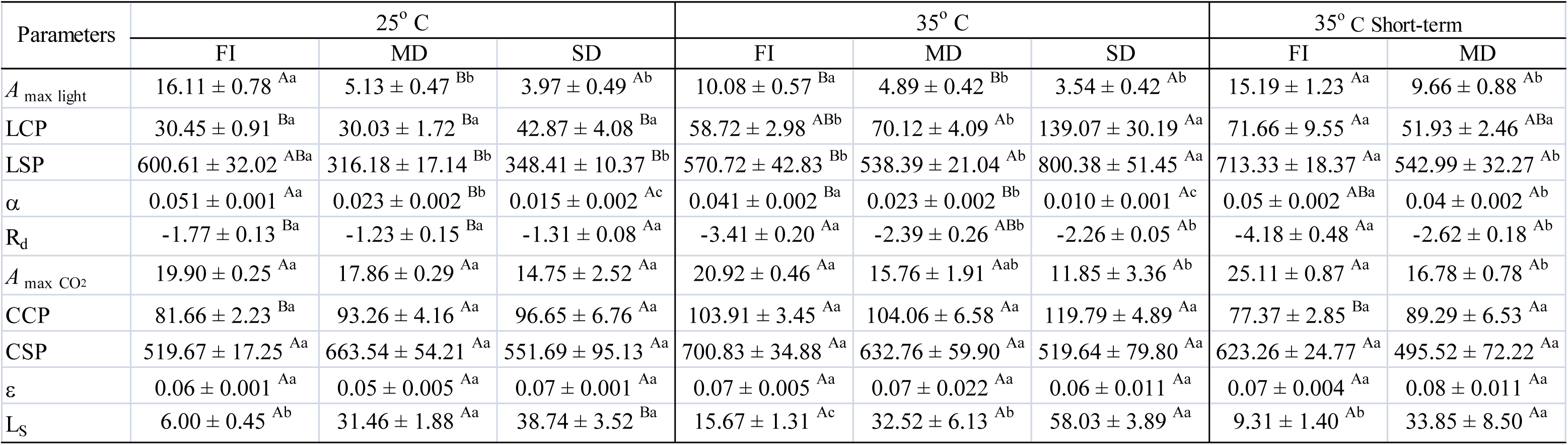
Parameters extracted from *A*_N_–PPFD and *A*_N_–C_i_ curves performed in leaves of *Eucalyptus globulus* grown at 25° C, 35° C and 35° C Short–term under: full irrigation (FI), moderate drought (MD) and severe drought (SD). Data represent means ± SE (n=6). Different capital letters express significant differences among thermal treatments and different lower-case letters express significant differences among irrigation treatments (Tukey’s test, *p* < 0.05). *A*_max light_ = maximum CO_2_ assimilation rate measured in the light response curves (μmol CO_2_ m^-2^ s^-1^), LCP = light compensation point (μmol photons m^-2^ s^-1^), LSP = light saturation point (μmol photons m^-2^ s^-1^), α = apparent quantum efficiency, R_d_ = dark respiration (μmol CO_2_ m^-2^ s^-1^), *A*_max CO_2 = maximum CO_2_ assimilation rate measured in the CO_2_ response curves (μmol CO_2_ m^-2^ s^-1^), CCP = CO_2_ compensation point (μmol CO_2_ m^-2^ s^-1^), CSP = CO_2_ saturation point (μmol CO_2_ m^-2^ s^-1^), χ = apparent carboxylation efficiency, L_s_ = Stomatal limitation of photosynthesis (%).

As with CO_2_ assimilation rates and g_s_, irrigation treatments affected intrinsic water use efficiency (WUE_i_), showing significantly higher values under water deficit in all thermal treatments (Figure 3). Despite of the non–significant differences in FI among the thermal treatments, there was a trend to increase in WUE_i_ under water deficit at 35° C which is not pronounced at 35° C Short– term, suggesting that the stomatal control under drought is influenced by the exposure time to 35° C. Dark respiration (R_d_) increased with the temperature and decreased with the water deficit in all irrigation treatments (Table 2).

Parameters derived from *A*_N_–PPFD curves showed marked differences among the thermal treatments (Table 2). Light compensation point (LCP) was significantly higher in all irrigation treatments at 35° C and 35° C Short–term compared to 25° C. Moreover, while LCP increased under MD and SD at 35° C when compared to FI, at 35° C Short–term it decreased. Light saturation point (LSP) decreased significantly with water deficit at 25° C and 35° C Short–term, while at 35° C it was significantly higher under SD. The apparent quantum efficiency (α) showed a similar response in all thermal treatments, consisting in a significant decrease with the increase of water deficit severity. However, there was a lower decrease in α under a short–term exposure to 35° C, which showed the highest values under MD compared to other thermal treatments.

Parameters derived from CO_2_ response curves (*A*_N_–C_i_) were mostly affected by water deficit (Table 2). Maximum assimilation rate (*A*_max CO_2) decreased with water deficit but no significant difference was detected among thermal treatments. Whereas the CO_2_ compensation point (CCP) increased with water deficit in all thermal treatments, showing a trend to be higher at 35° C, CO_2_ saturation point (CSP) did not show a consistent trend at 25° C, while at 35° C and 35° C Short– term it decreased with water deficit severity. The apparent carboxilation efficiency (ε) did not show a clear variation pattern comparing irrigation treatments in all thermal treatments studied (Table 2).

The relationship between net CO_2_ assimilation (*A*_N_) and intercellular CO_2_ concentration (C_i_), mesophyll conductance (*g*_m_) and stomatal conductance (*g*_s_), maximum electron transport rate (*J*_max_) and Rubisco maximum carboxylation rate (*V*_cmax_) extracted from *A*_N_–C_i_ curves are presented in Figure 4, and such parameters showed distinct responses to water deficit in the different thermal treatments. The decrease in *A*_N_ with water deficit at 25° C and 35° C is probably related with the decrease in *g*_s_ and *g*_m_, which caused a substantial decrease in C_i_ in the leaf mesophyll, together with the decrease in *V*_cmax_ and *J*_max_ which occurred mainly at 35° C. However, the high *A*_N_ rates observed at 35° C Short–term is probably related with the maintenance of a high *g*_m_, which would have supported a high C_i_ even under a low *g*_s_. Moreover, the maintenance of a high *V*_cmax_ and *J*_max_ can also have contributed to the high *A*_N_ at 35° C Short–term (Figure 4).

**Figure 4.**
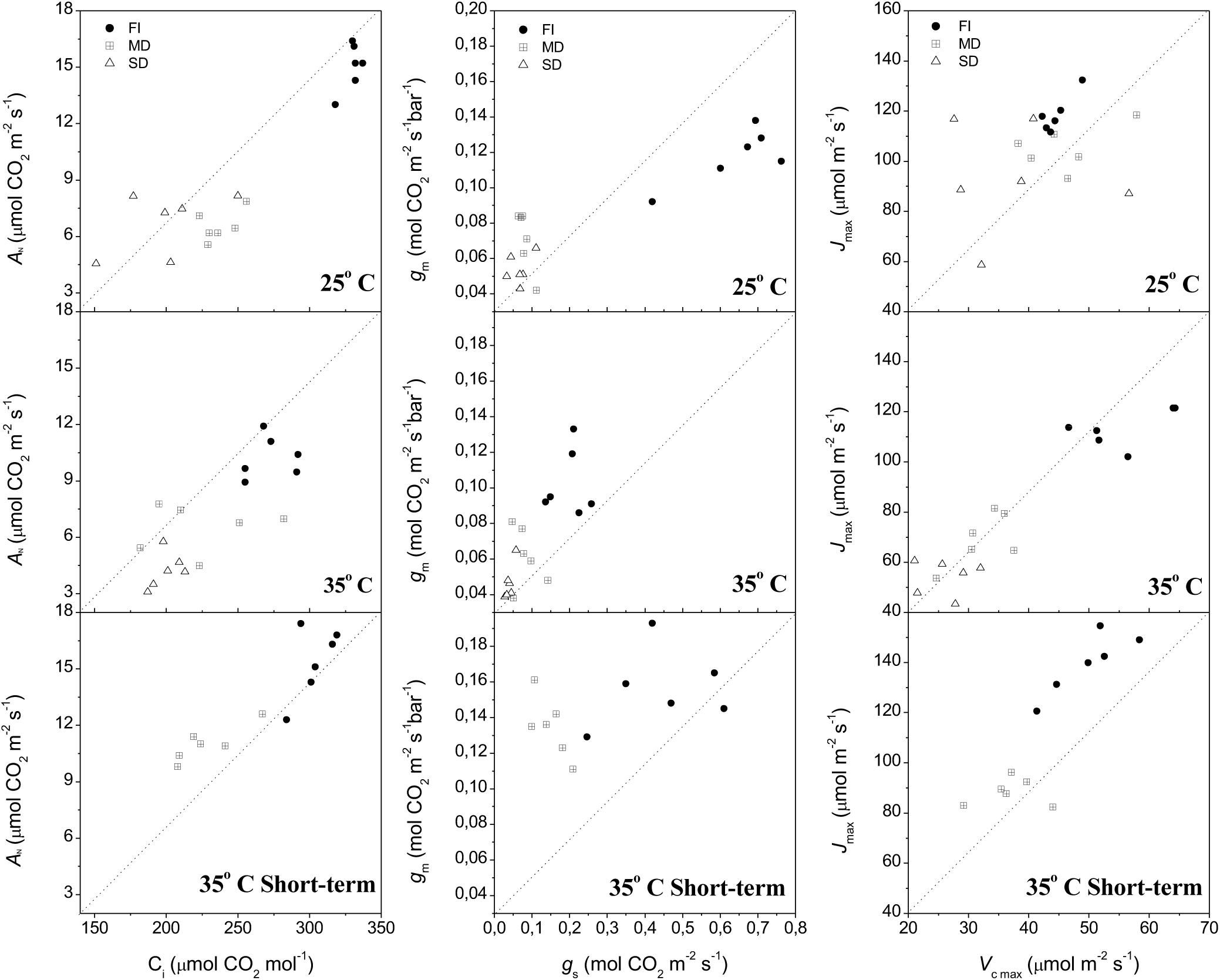
Relationship between net CO_2_ assimilation (*A*_N_) and intercellular CO_2_ concentration (C_i_), mesophyll (*g*_m_) and stomatal conductance (*g*_s_), maximum electron transport rate (*J*_max_) and maximum rate of Rubisco carboxylation (*V*_cmax_) extracted from CO_2_ response curves at 380 μmol CO_2_ m^-2^ s^-1^, performed in leaves of *Eucalyptus globulus* grown at 25° C, 35° C and 35° C Short-term under different irrigation treatments: full irrigation (FI), moderate drought (MD) and severe drought (SD). Data represent individual measurements (n=6).

In agreement with the progressive decrease in *g*_s_ with water deficit increase, the stomatal limitation of photosynthesis (L_s_) increased significantly in response to drought in all thermal treatments studied, being even higher at 35° C which reached more than 3–fold higher values under SD compared to FI (Table 2).

The principal component analyses (PCA) graph shows the representation of all thermal and irrigation treatments on the two major principal components, presenting some degree of separation among them (Figure 5). The first principal component (PC1) accounted for 49.3% of total data variance, while the second principal component (PC2) accounted for 16.8%, jointly explaining 66.1% of the data. Irrigation regimes presented a clear separation within thermal treatments, mainly along PC1, where the most significant variables composing the first principal, in order of importance, were: α, *A*_N_, *A*_max light_, L_s_ and *J*_max_. Thermal treatments were most distinguishable along PC2, where the most significant photosynthetic parameters were: LSP, LCP, F_V_/F_0_, F_V_/F_M_ and R_d_.

**Figure 5.**
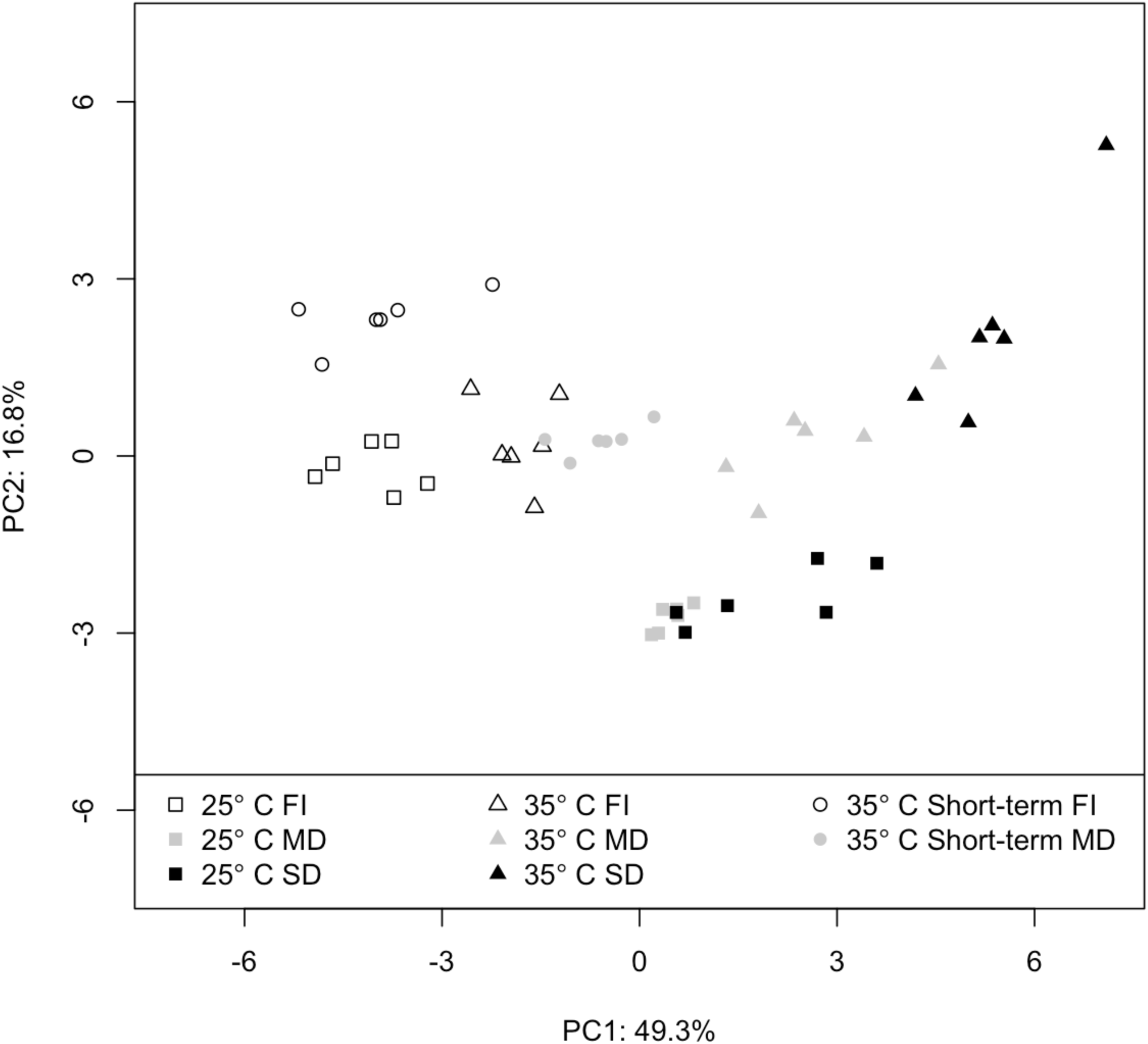
Distribution of water regimes in the thermal treatments for first and second principal components, PC1 and PC2 axes, respectively.

## DISCUSSION

### Effects on photochemical apparatus

It has been reported that the photochemical apparatus presents high robustness and resilience to drought (CORNIC et al., 1989; EPRON & DREYER, 1992, 1993). In fact, our results demonstrated that photochemical efficiency of PSII was not markedly sensitive to water deficit, mainly in plants at 25° C and 35° C Short–term (Table 1). Chlorophyll fluorescence parameters evaluated in dark–adapted leaves of *E. globulus* indicated that the photochemical apparatus was more affected by water deficit when plants were exposed to long–term 35° C, since a substantial increase in F_0_ and F_M_ and decrease in F_V_/F_M_ and F_V_/F_0_ were observed. This is compatible with results shown by HAVAUX (1992) which reported that exposure of potato leaves to high temperatures caused an increase in PSII activity in water–stressed plants. A prolonged exposition to 35° C caused a higher damage to the photochemical apparatus in *E. globulus*, since PSII is highly thermolabile and its activity is greatly reduced or even partially stopped under high temperatures (BUKHOV et al., 1999; CAMEJO et al., 2005), mainly due its localization in the thylakoid membranes which generally are damaged by heat. Moreover, the chlorophyll fluorescence yield in the dark–adapted state indicated that the photochemical apparatus was apparently more sensitive to temperature than to water deficit (compare values under FI between 25° C and 35° C with differences between 25° C FI, MD and SD in Table 1).

The decrease in F_V_/F_M_ coupled to an increase in F_0_ indicates that plants grown at 35° C under MD and SD conditions suffered damage in the photochemical apparatus and, consequently, presented photoinhibition in response to water deficit. This suggests that photoinhibition only occurs after a long–term exposure to 35° C in conjunction with water deficit, once plants of *E. globulus* grown at 35° C Short–term under drought showed no signs of photochemical damage. Thus, the absence of a substantial reduction in the efficiency of photosystem II indicates a considerable stability of this machinery in *E. globulus*, mainly in short–term exposure to moderately high temperatures. Moreover, the short–term exposure to 35° C apparently induced a protective mechanism in plants under water deficit, since it was observed a decrease in F_0_ while F_M_ was practically unchanged, and F_V_/F_M_ and F_V_/F_0_ increased slightly under MD when compared do FI (Table 1).

In contrast to F_V_/F_M_, ΦPSII and qP indicated that *E. globulus* was more sensitive to water deficit than to heat stress since the courses of both parameters were lower in plants under water deficit, while remaining practically unchanged with the temperature increase in well–watered plants (Figure 2). Interestingly, we observed that water deficit had a more pronounced effect in chlorophyll fluorescence parameters measured in light rather than in dark–adapted leaves, which showed more sensibility to temperature increase. Possibly, this difference reflects the fact that dark– adapted leaves have closed stomata implying in less means to dissipate energy and probably resulting in greater temperature sensitivity. This could superimpose the effect of water deficit, which apparently acts mainly through stomatal closure.

The suppression of NPQ due to temperature stress, observed between 25° C FI and 35° C FI (Figure 2), has been previously reported in barley by KALITUHO et al. (2003). NPQ increase under water deficit, observed in all thermal treatments but significantly less in both 35 ° C treatments, has been associated with a photoprotective role by avoiding excessive energy influx into the photosystem (OSMOND, 1994). Thus, apparently the short–term exposure to 35 ° C did not cause significant damage since there was no reduction in NPQ, whereas the response to water deficit could have been compromise by both short and long–term treatments. Furthermore, the indications of photoinhibition at 35° C under MD and SD conditions may be related with limitations in the NPQ, since damage to PSII is attenuated by NPQ or locally by cyclic electron transport (RUMEAU et al., 2007). Variations in ETR in all irrigation and thermal treatments followed similar trends to *A*_N_, probably due to the fact that Rubisco activity is highly correlated with ETR (KRALL et al., 1995). Previous studies showed that heat shock alters the photosynthetic activity via suppression of chloroplast electron transport (PASTENES & HORTON, 1996; FELLER et al., 1998), which is compatible with the ETR reduction observed at 35° C, possibly also related with an increase in the Rubisco oxygenase activity leading to higher photorespiration. However, at 35° C Short–term, there was no decrease in ETR under FI while there were higher values and a lower reduction of ETR under MD when compared with other thermal treatments, suggesting an increase in tolerance to water deficit.

As proposed by hypothesis *i*, which states that *E. globulus* show different responses to short and long–term exposure to 35° C, the chlorophyll fluorescence parameters indicated that the photochemical apparatus was differently affected by the time of exposure to 35° C. Whereas the long–term exposition compromised more the photochemical functions, the short–term treatment even had a potential protective effect, ensuring the photochemical efficiency and increasing the tolerance to water deficit. Hypothesis *ii*, stating the effect of both stresses combined will not be simply the sum of each condition applied separately, was also supported since the combined effect of temperature and water deficit revealed to be synergistic in some cases, especially through chlorophyll fluorescence parameters measured in dark–adapted leaves at 35° C, and even enhancing tolerance mechanisms, as indicated by the 35° C Short–term treatment.

### Effects on carbon assimilation

The effects of water deficit on photosynthesis were widely described by several authors (CHAVES, 1991; CHAVES et al., 2003; CORNIC, 2000, FLEXAS et al., 2004; CHAVES et al., 2009; LAWLOR & TEZARA, 2009) and there is a general consensus that the decrease in the photosynthetic rate is caused primarily by stomatal closure, which limits the availability of CO_2_ within the leaf. As expected, we verified a substantial decrease in *g*_s_ in plants under water deficit, in all thermal treatments studied, with a proportional decrease in C_i_ (Figure 3, 4). Interestingly, a short–term exposure to 35° C did not cause a substantial stomatal closure under FI, and furthermore resulted in a lower reduction in *g*_s_ in response to water deficit, allowing the maintenance of high *A*_N_ rates compared to 25° C and 35° C. Possibly, the short–term exposure to 35° C induces a protective mechanism which allows the maintenance of a high *g*_s_, increasing the tolerance to water deficit.

Moderately high temperatures increased stomatal limitation (L_s_) both under FI and water deficit (Table 2). Moreover, long term exposure to 35 ° C induced a substantial decrease in *g*_s_ in well–water plants, probably causing the observed reduction in *A*_N_ and C_i_ (Figure 3). Such decrease in *A*_N_ under FI at 35° C indicates a down–acclimation of well–watered plants when exposed to moderately high temperatures for a long time. Moderate heat stress reduces *A*_N_ and *g*_s_ in many plant species due to decreases in the activation state of Rubisco (CRAFTS–BRANDNER & SALVUCCI, 2002), once Rubisco activase is thermolabile (FELLER et al., 1998; SALVUCCI et al., 2001). These authors propose that the heat–induced deactivation of Rubisco is the primary constraint on photosynthesis at moderately high temperatures, showing that chlorophyll fluorescence signals from PSII are not affected by the same temperatures (CRAFTS–BRANDNER & SALVUCCI, 2000; HALDIMANN & FELLER, 2004) and injuries or death may occur only after long–term exposure. Accordingly, we verified that the thermal treatments caused more pronounced effects in the carbon assimilation than in the photochemical apparatus, and injuries occurred mainly after a long–term exposure to high temperature (Figure 2, 3).

The reduction in *g*_m_ is another component that may influence carbon availability to photosynthesis under stress conditions, being regulated almost simultaneously to *g*_s_ (FLEXAS et al., 2008). LAISK et al. (1998) proposed that the decrease in *A*_N_ in response to temperature might be partially related with *g*_m_, while several authors reported *g*_m_ reduction under water stress (SCARTAZZA et al., 1998; FLEXAS et al., 2002, 2008; GALMÉS et al., 2007). Accordingly, we also observed a decrease in *g*_m_ in response to water deficit and temperature increase, possibly underlying the decrease in *A*_N_ (Figure 4). However, the short–term exposure to 35° C induced the maintenance of a high *g*_m_ under water deficit, which probably enabled a high C_i_ even with the low *g*_s_ thus contributing to the high *A*_N_ under MD (Figure 4). Jointly, the maintenance of high *V*_cmax_ and *J*_max_ rates could also have contributed to the high *A*_N_ rates under MD at 35° C Short–term, since under long–term exposure to 35° C there was a more pronounced decrease in *V*_cmax_ and *J*_max_ under drought. Previous studies have found that the *J*_max_:*V*_cmax_ ratio tends to decrease with increasing growth temperature (BUNCE, 2000; KATTGE & KNORR, 2007). However, the short-term exposure to 35° C slightly increased this ratio, showing the highest *J*_max_:*V*_cmax_ under MD, also contributing to the maintenance of a high *A*_N_ under water deficit (Table 1).

The increase in R_d_ is another factor that contributed to the decrease in CO_2_ assimilation rate (Table 2). WAHID et al. (2007) reported that elevated temperature causes a heat–induced imbalance in photosynthesis and respiration since, in general, photosynthetic rate decreases while dark and photo–respiration increase considerably under high temperatures. Correspondingly, we observed a substantial increase in R_d_ in short and long–term exposure to 35° C under FI, while plants under water deficit showed a decrease in R_d_ in all thermal treatments. This response may save energy under stress conditions. However, higher R_d_ did not correlate directly with photosynthetic performance given that plants at 35° C Short–term showed the highest R_d_ rate in both FI and MD, but maintained equal or higher *A*_N_ than the remaining thermal treatments under all irrigation conditions. Decreases in R_d_ with water deficit observed at 25° C and 35° C did not lower the LCP, probably because the apparent quantum efficiency (α) had a significant decreasing trend with drought severity. This was not the case at 35° C Short–term which showed the lowest reduction in α and a decrease in LCP under water deficit, having significantly higher α than the remaining thermal treatments under MD. In fact, although *A*_max light_ decreased with water deficit in all thermal treatments, the lowest reduction occurred at 35° C Short–term which presented the highest maximum assimilation rate under MD, being a further indicative of the increase in tolerance to water deficit.

Carbon assimilation evaluated through gas–exchange parameters showed that plants respond differently to water deficit according to the thermal treatment. A general response to drought is the reduction in *g*_s_ and *g*_m_, which implies in a reduction of CO_2_ diffusion in the mesophyll consequently decreasing the photosynthetic activity, as evidenced by decreases in α and *A*_max_ (Table 2). Moreover, *V*_cmax_ and *J*_max_ also decreased substantially with water deficit, mainly under long–term exposure to 35° C. The combination of water deficit and a long–term exposure to 35° C caused a down–acclimation and had the most negative effect in photosynthesis, while a short–term exposure to 35° C implied in a less compromised photosynthetic apparatus having, a higher performance under drought even when compared to the control 25° C condition, supporting hypothesis *i* and *ii*. Possibly, the main factor accounting for the higher photosynthetic efficiency of plants exposed to 35° C Short–term under water deficit is the maintenance of a relatively high stomatal and mesophyll conductance, showing an especially lower reduction in *g*_m_ in response to drought (Figure 4). Since the response to a combination of heat and drought differed from the superposition of individual responses, as observed mainly at 35° C, the hypothesis *ii* was also confirmed by our results.

The combination of PC1, capturing 49.3% of the variance contained in the data, with PC2, 16.8%, yielded a considerably coherent clusters of the replicates according to irrigation and thermal treatments (Figure 5). There was a clear gradient of irrigation treatments along the main axis of variation, indicating a consistent overall effect of water deficit in the global change of physiological variables. Conversely, PC2 mainly accounted for the separation among thermal treatments, which are less discernible along PC1. This is in general agreement with the statistical results obtained with the Tukey’s test, since the main parameters composing PC1 (α, *A*_N_, *A*_max light_, L_s_ and *J*_max_) differed more with respect to irrigation, whereas the parameters composing PC2 (LSP, LCP, F_V_/F_0_, F_V_/F_M_ and R_d_) differed more according to thermal treatment.

## CONCLUSIONS

The co–occurrence of water deficit and moderately high temperature in *E. globulus* amplified the reduction of photosynthetic rates, compared to the single effects, mainly after a long– term exposition to 35° C. This effect was less pronounced in the exposure to 35° C Short–term, which apparently increased the tolerance to water deficit given the observed highest overall photosynthetic efficiency under drought. When compared to other thermal treatments under water deficit, the short–term exposure to 35° C induced higher *A*_N_, *A*_max light_, α, ETR, *V*_cmax_, *J*_max_, *g*_m_, *g*_s_ and C_i_, lower F_0_ and LCP, lower reduction in <!PSII and qP and no reduction in F_M_, F_V_/F_M_ and F_V_/F_0_, suggesting an enhancement of photosynthetic capacity under water deficit, possibly through a cross–tolerance mechanism.

Hypothesis *i* was supported by the results presented herein since the effects of short and long–term exposure to 35° C differed, being a longer exposure generally more injurious to the photosynthetic efficiency. Furthermore, the short–term exposure to 35° C showed a fortunate protective effect under water deficit, apparently as a result of maintaining a high stomatal and mesophyll conductance. Hypothesis *ii* was also supported by the results since water deficit and moderately high temperature had a singular effect when applied in combination, distinct from the mere sum of their individual effects. This is in accordance to MITTLER (2006) who proposes that a particular stress combination should be regarded as a singular state of abiotic stress requiring a new defense or acclimation response.

Our results allowed to evaluate the individual and combined effects of water deficit and different times of exposure to a moderately high temperature, which is of crucial importance given that both stresses often co–occur in the field. This is particularly relevant given the lack of knowledge regarding physiological responses to combined stress types, especially in *Eucalyptus*, and the imminent global climate change which will likely increase the incidence of drought and moderately high temperature. Thus, the present study showed that the acclimation to a long–term exposure to 35° C impose some restrictions to photosynthetic activity mainly under water deficit, while a short–term exposure enhances the photosynthetic efficiency under water deficit, conferring some degree of tolerance. Beyond contributing to the understanding of *E. globulus* physiological responses to realistic stress conditions, we hope to assert the importance of evaluating the effects of stress combinations which can reveal non expected responses, while potentially providing insights into underlying physiological mechanisms involved in acclimation.

